# Potential Immunomodulatory Role of Lead in Monocyte/Macrophage Differentiation

**DOI:** 10.1101/2023.04.03.535415

**Authors:** Stacia M. Nicholson, Francis A.X. Schanne

**Affiliations:** St. John’s University, Department of Pharmaceutical Science, New York, 11439, USA; Columbia University Mailman School of Public Health, Department of Environmental Health Sciences, New York, 10032, USA

## Abstract

Lead (Pb) is a pernicious toxic metal and public health hazard, as it persists in the environment contaminating soil, food, and drinking water. Lead exerts its toxic effect on numerous organ systems, with the abundance of focus on the red blood cells and neurons of the hematopoietic and central nervous systems. However, insufficient investigation has been conducted on the effect of Pb on immune cells. In the current study, the toxic effects of Pb on immune cells of monocyte/macrophage lineage are described. Pb exerts a dose-dependent alteration in differentiation of monocyte/macrophage cells that retain some plasticity in development. Pb induces a bifurcation in differentiation of monocyte/macrophage cells, resulting in inhibition of osteoclastogenesis and induction of dendritic cells (DC). This phenomenon was demonstrated in RAW 264.7 murine monocyte/macrophage cell line and was consistent with response in rat bone-marrow derived macrophage (Sprague-Dawley). Pb primarily produced this response through induction of GM-CSF production and inhibition of p38/MAPK activity. Long-term exposure to physiologically relevant concentrations of Pb has the potential to modulate the immune system through altering the cell-lineage commitment of monocyte/macrophage lineage cells in a dose-dependent manner. Thus, Pb may function as an immunomodulator.

## Introduction

Lead is a pernicious toxic metal that persists in the environment despite being banned from use in gasoline, paint, and pesticides in the United States by the late 1980’s (1-3). Lead exposure occurs through inhalation and ingestion of contaminated food, water, and air, with adults and children continuing to be exposed to Pb from dust, paint chips, and lead pipes found in old home and infrastructure (4, 5). In addition, growing sources of Pb exposure in adults occur through substance abuse (cigarette smoking and opium), traditional or herbal medicines, and gunshot wounds (6, 7). Pb is toxic to several organ systems (8), however, most attention has been given to its neurotoxic and hemotoxic effects which most profoundly affect children resulting in neurological disorders, learning and behavioral problems, and anemia (9-11). Pb is absorbed via the lung and gastrointestinal system into the blood, where it is circulated to the soft tissues before being sequestered in bone (12). Pb is removed from the body via biliary and urinary excretion, having a half-life of 35 and 40 days in the blood and soft tissue, respectively (13, 14). However, in the bone Pb has a half-life of 2 to 3 decades, and can cycle back into the blood during growth, injury, pregnancy, lactation, menopause, or any remodeling of the skeletal system (14-16).

Through environmental exposure and endogenous cycling of Pb from bone into blood, exposure to Pb is continuous, but most *in vivo* and *in vitro* studies have looked at the acute toxic effects of Pb exposure. Pb has been reported to cause osteopenia (17) and osteopetrosis, but data on osteoporosis has been inconsistent. However, Pb’s effect on the monocyte/macrophage cells of the hematopoietic system has not been of significant focus, though monocyte/macrophage cells have their origins in the bone-marrow, and are implicated in the pathology of various immune related diseases throughout the body. When the role of Pb on the immune response of monocyte/macrophage cells has been investigated, most studies involved the use of LPS to simulate infection or inflammation through TNF-alpha (TNF-α) induction (18-24).

Typical *in vitro* toxicity methods involve exposing cells for a period ranging from 6 to 72 h, which represents acute toxicity (25). However, because humans are consistently exposed to low levels of Pb, through the environment, we wanted to determine the chronic toxic effects of physiologically relevant levels of Pb.

Therefore, we investigated the effect of long-term in vitro exposure (prolonged or continuous; 1 ≤ 5 to 9 d) to Pb in monocyte/macrophage cells in the absence of inflammation. Using RAW 264.7 murine cell-line preliminary experiments were carried out to determine dose-response and acute effect on cell viability which demonstrated cell survival. With prolonged exposure times, observational analysis by microscopy revealed proliferative and phenotypic changes (size, morphology, adherence) indicative of cellular differentiation and dendritic cell (DC) formation. Pb was previously described as having the ability to induce the formation of pre-osteoclasts or osteoclast-like multinucleated cells in bone-marrow derived macrophage (26), and RAW 264.7 monocyte/macrophage cell line has also been used extensively in research as a model for osteoclastogenesis (27). Osteoclastogenesis is the process by which monocyte/macrophage cells differentiate into TRAP (tartrate-resistant acid phosphatase) producing giant cells, known as osteoclasts, a type of multinucleated macrophage. RAW 264.7 cells being macrophages express the M-CSF (macrophage colony stimulating factor) receptor and produce soluble M-CSF through the shedding of this receptor (28), and therefore can be differentiated in the presence of Receptor-Activator of Nuclear-factor Kappa-b Ligand (RANKL) to form osteoclasts. M-CSF and RANKL are essential for osteoclastogenesis (29). As such, we used RAW 264.7 cell induction to osteoclast as a model for studying Pbs effect on differentiation. Using this model, we examined the effect of Pb on differentiating monocyte/macrophage cells to monocyte-derived dendritic cells and osteoclasts. We determined that Pb may function as an immune modulator due to its ability to suppress the formation of multinucleated macrophage and drive dendritic cell (DC) induction.

Furthermore, experiments were replicated in bone-marrow cells, progenitors of bone-marrow derived monocyte/macrophage (BMMØ), to determine whether the effect on differentiation seen in mature monocyte/macrophage cells also occurred in the hematopoietic system from which macrophage arise. These exposures were carried out in rat primary cells to also demonstrate that the effects seen in RAW 264.7 cells were not owed to species or cell-line characteristics.

Moreover, to elucidate the mechanistic action of Pb on differentiation, pathways involved the process were examined. Previous studies by other labs have revealed the involvement of the PKC-MAPK pathway in acute Pb toxicity on monocyte/macrophage cells associated with an inflammatory response (21). However, in the current study, it was identified that with pro-longed exposure, Pb’s action on p38/MAPK activity is associated with halted proliferation and the induction of differentiation. These findings were accompanied by the reduction of inflammatory marker TNF-α, NFκB, and IL-6, which also play a role in the PKC-MAPK differentiation pathway.

## Methods & Materials

RAW 264.7 (TIB-71, ATCC) and RAW-Blue (Invivogen) cells were acquired from the Dr. Lin Mantell Lab at St. John’s University. Bone-marrow cells were isolated from the femurs and tibias of 4 – 6 w old Sprague-Dawley rats bred in-house (animal use protocol was reviewed and approved by St. John’s University’s institutional Animal Care and Use Committee (IUCAC)). and stimulated with M-CSF (macrophage colony stimulating factor) derived from LADMAC cells (CRL-2420, ATCC) to form bone-marrow derived macrophages (BMMØ).

Lead (II) acetate trihydrate, 99% (Alfa Aesar, England) was dissolved directing into complete media (Media + 10% FBS +1 % Penicillin/Streptomycin) and diluted to the concentrations of 0.01, 0.1, 1.0, 5.0, and 10 µM for all experiments. Each concentration was tested in triplicate for all experiments.

Recombinant mouse TNFSF11 (IBI Scientific, Indiana), also known as RANKL, was solubilized in sterile phosphate-buffered saline and 0.1% BSA. In all experiments where applicable, a final concentration of 30 ng/ml RANKL was used.

### Viability and Proliferation Assays for Baseline Acute Toxicity

Cell viability and proliferation were determined using trypan blue exclusion assay and 3- (4,5Dimethylthiazol-2-Yl)-2,5-Diphenyltetrazolium Bromide (MTT) proliferation assay, respectively. RAW 254.7 cells were seeded into 6-well tissue culture treated plates at a density of 500,000 cells per well, in DMEM complete growth media, and incubated overnight in humidified air at 37°C and 5 % CO2. Media was removed, and replaced with DMEM complete growth media containing Pb. Control cells did not receive Pb, but did receive fresh media. Cells were further incubated for 24 h. Cells were washed with PBS and collected via scraping. After addition of trypan blue, cells were observed under light microscopy and counted using a hemocytometer. Viability was determined by the number of live cells which excluded trypan blue versus the number of dead cells which took up the dye and were blue. Proliferation was assessed by comparing the number of cells present at the completion of the experiment versus the number of cells initially seeded; accounting for dead cells. MTT assay was carried out under the same treatments as trypan-blue exclusion assay to validate results. Pb was removed after 24 h exposure, and cells were allowed to proliferate for an additional 24 in incubation at 37°C and 5% CO2. MTT solution was then added directly into wells without removing media or washing. Cells were incubated with MTT solution for 30 min and converted dye was solubilized with DMSO. Optical density was measured at 540 nm on a microplate reader and proliferation was reported as a percent of the final cell density in each sample in comparison to the control. All exposures were carried out in triplicate.

### Long-term Pb Exposure & Multinucleated cell (osteoclast) Differentiation

RAW 264.7 cells were seeded at 200,000 cells/well in 6-well plates in DMEM complete growth media and incubated overnight in humidified air at 37°C and 5% CO2, to allow cells to attach.

Growth media was replaced with MEM complete media containing Pb alone or in combination with RANKL. Cells were then further incubated for 5 to 7 d. Supernatant was collected and used for the quantification of TRAP production via spectrophotometry. Cells were fixed with 4% paraformaldehyde (Alfa Aesar, Ward Hill, MA) and stained with the tartrate-resistant acid phosphatase (TRAP) detection solution for the identification of osteoclasts (27). Giant multinucleated cells that contained greater than 2 nuclei and stained positive for TRAP were considered osteoclasts. Images were acquired with microscope camera and software, Motic Images Plus 3.0. Osteoclasts were counted via light microscopy.

### *NFκB &* TRAP Quantification

#### RAW-Blue™

(Invivogen) cells are RAW 264.7 cells with chromosomal integration of a secreted embryonic alkaline phosphatase (SEAP) reporter construct inducible by NFκB activation. Cells were seeded at 250,000 cells per well in 6-well plates and allowed to attach overnight in humidified incubation at 37°C and 5% CO2. Pb in MEM complete media was added in concentrations ranging from 0.01 to 10 µM, with and without RANKL. Supernatant was collected at 24 h and 5 d. Colorimetric detection reagent was added to each sample and alkaline phosphatase production, representative of NFκB activity, was quantified via absorbance (620 nm) using a microplate reader.

#### IL-1α Production

Supernatant from RAW 264.7 cells exposed to Pb (ranging from 0.01 to 10 µM) alone or in conjunction with RANKL for 7 d was used for the detection of IL-1α by ELISA kit (ebioscience). Colorimetric results were read on a microplate reader at absorbance 450nm.

#### TNF-alpha and GM-CSF Production

RAW 264.7 cells were seeded 200,000 cells/well in 6-well plates in DMEM complete growth media, and incubated overnight in humidified air at 37°C and 5% CO2, to allow cells to attach. Growth media was removed and MEM complete media containing Pb only or Pb in combination with RANKL was added. Cells were returned to incubation for 5 d to 7 d. The supernatant was collected for quantification of TNF-α and GM-CSF production and assessed using mouse TNF-α ELISA kit (ebioscience) and mouse GM-CSF ELISA kit (ebioscience). Samples were diluted 1:2 for TNF-α ELISA, and colorimetric results were read on a microplate reader at absorbance 450nm.

#### p38/MAPK Activity

500,000 cells per well were seeded into 6 well plates in DMEM complete growth media and allowed to attach overnight; incubated in humidified air at 37°C and 5% CO2. Growth media was removed and cells were washed with PBS before addition of Pb in MEM complete media. To stop experiment, cells were placed in -20°C freezer for 5 to 10 min before lysis (cell were lysed directly in media). Activity was determined using MAPKp38 (Total) InstantOne ELISA kit (Invitrogen) and absorbance was read on a microplate reader.

#### CD11c and DEC205 Immunostaining

RAW 264.7 and BMMØ exposed to Pb ranging from 0.01 to 10 µM Pb for 5 to 9 d were collected and centrifuged at 2200 rpm for 5 min to remove media containing Pb. Cells were then washed with PBS and diluted to 1,000,000 cells per ml. Fc receptors were blocked for 30 min with Protein A (MP Biomedicals, OH) in Stain Buffer (554656, BD Pharmingen) containing sodium azide, to prevent non-specific binding. Immunofluorescence was performed with antibody markers for macrophage (CD11b) and dendritic cells (CD11c or DEC205) and analyzed via flow cytometry. Data was analyzed with IDEAS 6.2 software.

#### Phagocytosis

RAW 264.7 cells exposed to Pb only or Pb + RANKL for 5 to 7 days in MEM complete media were collected, counted, and reseeded at identical densities and allowed to attach overnight in incubation in humidified air at 37°C and 5% CO2 in MEM complete media. Cells were then washed and incubated in serum-free MEM media for 2 h prior to addition of 1.75-micron latex beads (Fluoresbrite Carboxylate YG Microspheres, Polysciences, Inc, PA). Cells were incubated with beads at 37°C for an additional 2 h. Phagocytosis was stopped by washing cells with cold PBS. Fluorescence of external beads that were not removed with washing, were quenched with 0.4% trypan blue and fluorescence was read on a microplate reader.

#### BM-derived monocyte/macrophage (BMMØ) Generation and Pb Exposure Isolation and Generation

Bone-marrow cells (BMC) were isolated from the limb long bones of 4- to 6-week-old Sprague-Dawley rats following the procedure described by (29). BMC were differentiated to BMMØ in MEM complete growth media containing M-CSF from LADMAC cell-line (CRL-2420, ATCC) conditioned media. LADMAC cells are a transformed cell-line from mouse bone-marrow cells, which produce M-CSF; capable of supporting *in vitro* proliferation of bone-marrow macrophage (31). Cells were seeded onto non-tissue culture treated petri dishes. BMMØ purity was determined by CD206 antibody staining.

#### Pb Exposure

BMMØ were exposed to Pb in MEM complete media for 5 to 7 days on tissue-culture treated 6 – well plates, and incubated in humidified air at 37°C and 5 % CO2. Cells were either fixed in 4 % paraformaldehyde and stained for TRAP or harvested for CD11c immunofluorescence staining.

#### Statistical Analysis

Statistical analysis was performed using GraphPad Prism 7.04 software. Statistical significance was determined using unpaired sample t tests and one-way ANOVA. P value less than 0.05 was considered significant. *, **, *** denotes significance of p < 0.05, 0.01, 0.001 respective

## Results

### Viability and Proliferation

Viability in RAW 264.7 cells exposed to Pb ranging from 1 to 10 µM for 24 h was examined using trypan blue exclusion. RAW 264.7 cells double every 24 h. Doubling rate decreased with exposure to concentrations of 1.0 – 10 µM Pb. Though the total number of live cells present decreased between 1 and 10 µM Pb (27% and 50 % fewer cells, respectively); there was no statistical difference in the number of dead cells across all concentrations compared to control (Figure 1A). Results indicate that Pb does NOT induce cell death in monocyte/macrophage cells, but inhibits proliferation. MTT proliferation assay was carried to validate whether Pb inhibited proliferation. Pb induced a 20 to 80 % reduction in proliferation, with significant inhibition occurring at 1, 5, and 10 µM (Figure 1B).

**Figure 1.**
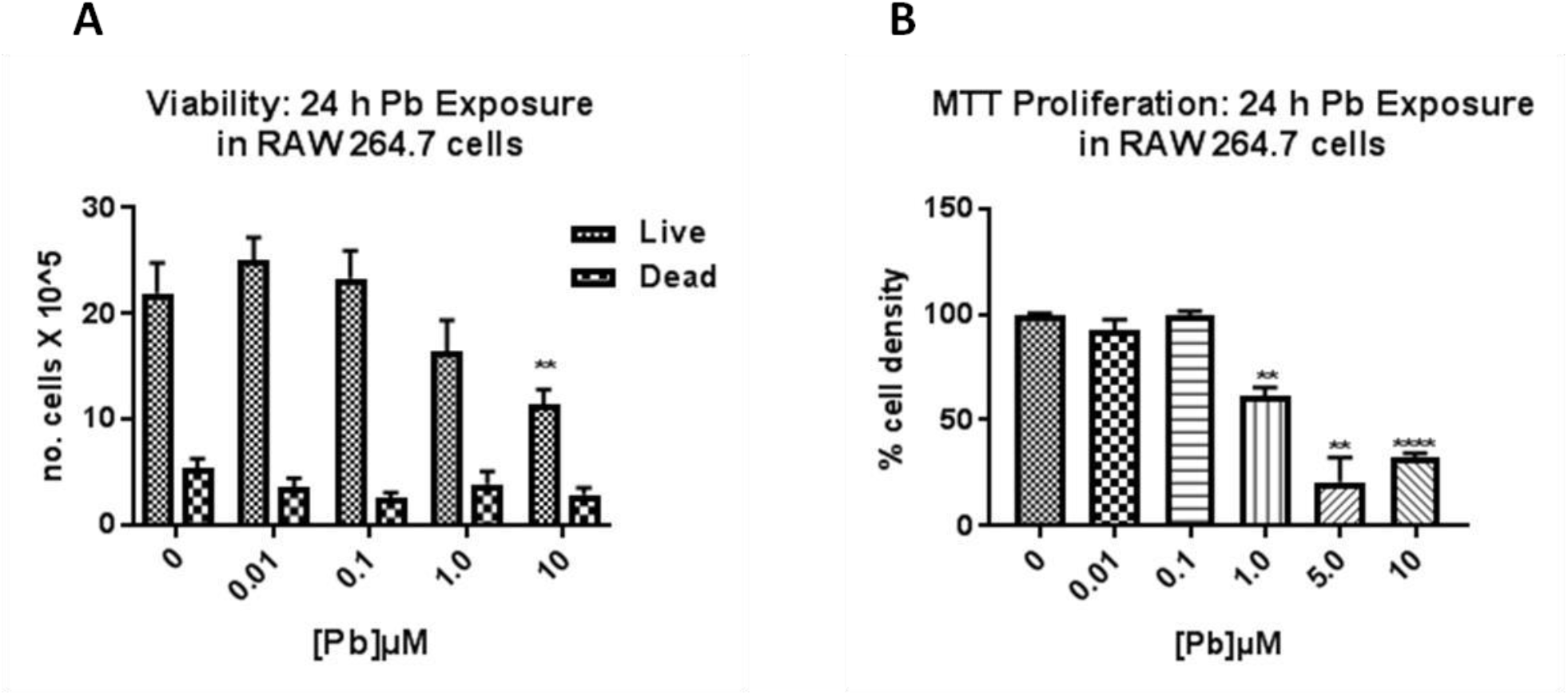
(A) Viability of RAW 264.7 cells exposed to Pb for 24 h (n = 3). (B) Proliferation of RAW 264.7 cells exposed to Pb 24 h (n = 3). The mean of each column was compared to the mean of the corresponding control. Unpaired *t* test was performed to determine statistical significance. **p ≤ 0.01, ****p≤ 0.0001.

### Pb Induced Differentiation

When long-term Pb exposures were carried out, RAW 264.7 cells exposed to 1.0, 5.0, and 10 µM Pb became amoeboid and non-adherent between 48 and 72 h, while cells in the control (0 µM), 0.01 µM and 0.1 µM exposures remained adherent (Figure 2A). Non-adherent cells appeared larger than control cells, and those exposed to 1.0 µM were single ameboid cell, but 5.0 and 10 µM cells formed primarily three-dimensional clusters of cells resembling raspberries or bunches of grapes (Figure 2B). These cell clusters appeared to have hair-like protrusions, and we referred to them as dendritic-clusters. It is important to note that control cells remained adherent even at full confluence indicating that non-adherence was not due to overgrowth.

**Figure 2.**
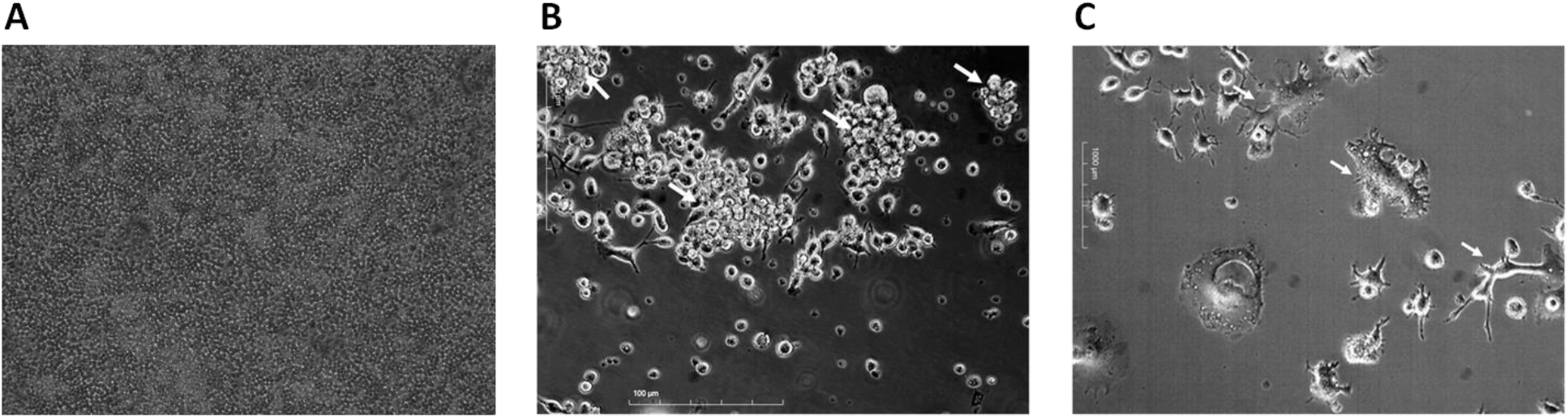
(A) Control RAW 264.7 cells at 100% confluency (B) Dendritic-clusters (5µM Pb). (C) Adherent Dendritic-cells after Pb wash-out (10 µM Pb). Images are representative. Images were captured via phase-contrast light microscopy at 20X magnification.

Furthermore, when dendritic-clusters were sub-cultured in Pb-free growth media, they separated and became adhered to the surface of the well. These cells varied in shape and size, but were larger than untreated RAW 264.7 cells, and displayed numerous dendritic processes indicative of DC (Figure 2C). These results suggest that prolonged Pb exposure induces differentiation in monocyte/macrophage cells.

### TNF-alpha Production

TNF-α is involved with cell proliferation and differentiation (32). The appearance of differentiated cells coincided with reduction in proliferation, pointing to a possible involvement of TNF-α. TNF-α production as assessed by ELISA showed a Pb induced dose-dependent decrease in the detection of the soluble cytokine in supernatant of cells exposed to Pb both acutely for 24 h and long-term for 5 d. Acute Pb exposure resulted in a 30 to 70 % reduction in TNF-α production, with the greatest inhibition occurring at 5 µM (Figure 3A). Long-term Pb exposure induced a 50 to 96 % reduction in TNF-α production in RAW 264.7 cells between 1.0 to 10 µM (Figure 3B).

**Figure 3.**
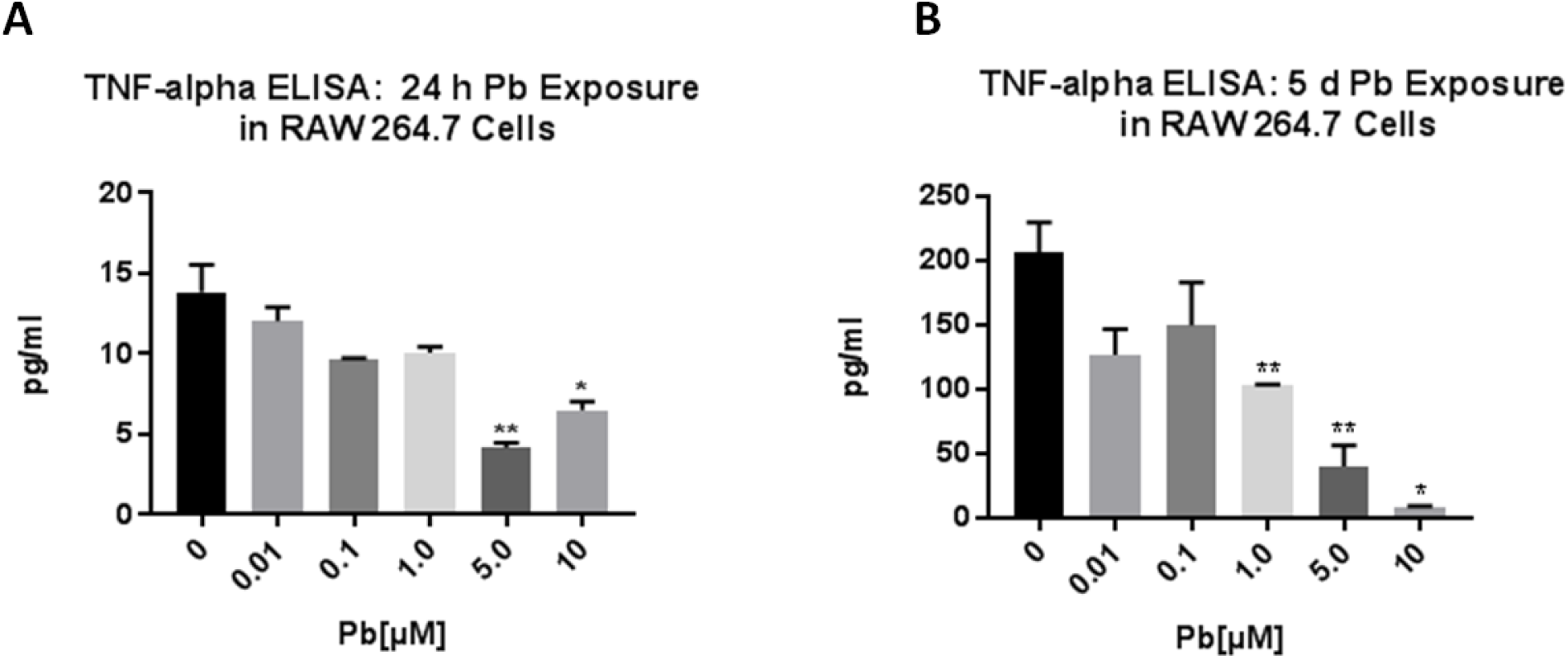
TNF-α production in RAW 264.7 cells, 24 h and 5 d Pb exposure (n =3). Pb dose-dependently decreased TNF-α production in RAW 264.7 cells exposed to Pb for 24 h at 5.0 and 10 µM, and 7 d at 1.0 to 10 µM Pb.

Dose-response decreased TNF-α levels also indicate that ameboid characteristic of cells seen primarily with 1.0 µM prolonged exposure were likely not due to inflammation induced activation.

### RANKL Stimulated Differentiation

Though Pb dose-dependently inhibited TNF-α production, it remained unclear what involvement this had with the induction of dendritic cell formation. TNF-α is known to induce differentiation of osteoclasts and to co-operate with RANKL in osteoclastogenesis (32). In addition, RAW 264.7 cells are a well characterized monocyte/macrophage cell line, which are routinely used for the generation of osteoclasts *in vitro* (27). Therefore, the effect of long-term Pb exposure on differentiation in monocyte/macrophage cells was studied using RANKL stimulation of RAW 264.7 cells as a model for osteoclastogenesis, to observe whether Pb would modulate this form of differentiation Long-term Pb exposure in RAW 264.7 cells stimulated with RANKL showed an over-all dose-dependent decrease in osteoclastogenesis. The number of osteoclasts formed decreased as the concentration of Pb increased, with significant inhibition occurring at concentrations of Pb ranging from 1.0 to 10 µM (Figure 4). Interestingly, non-adherent cells and dendritic clusters continued to form under stimulation of osteoclastogenesis by RANKL. The inhibition of osteoclastogenesis was reciprocal to DC formation.

**Figure 4.**
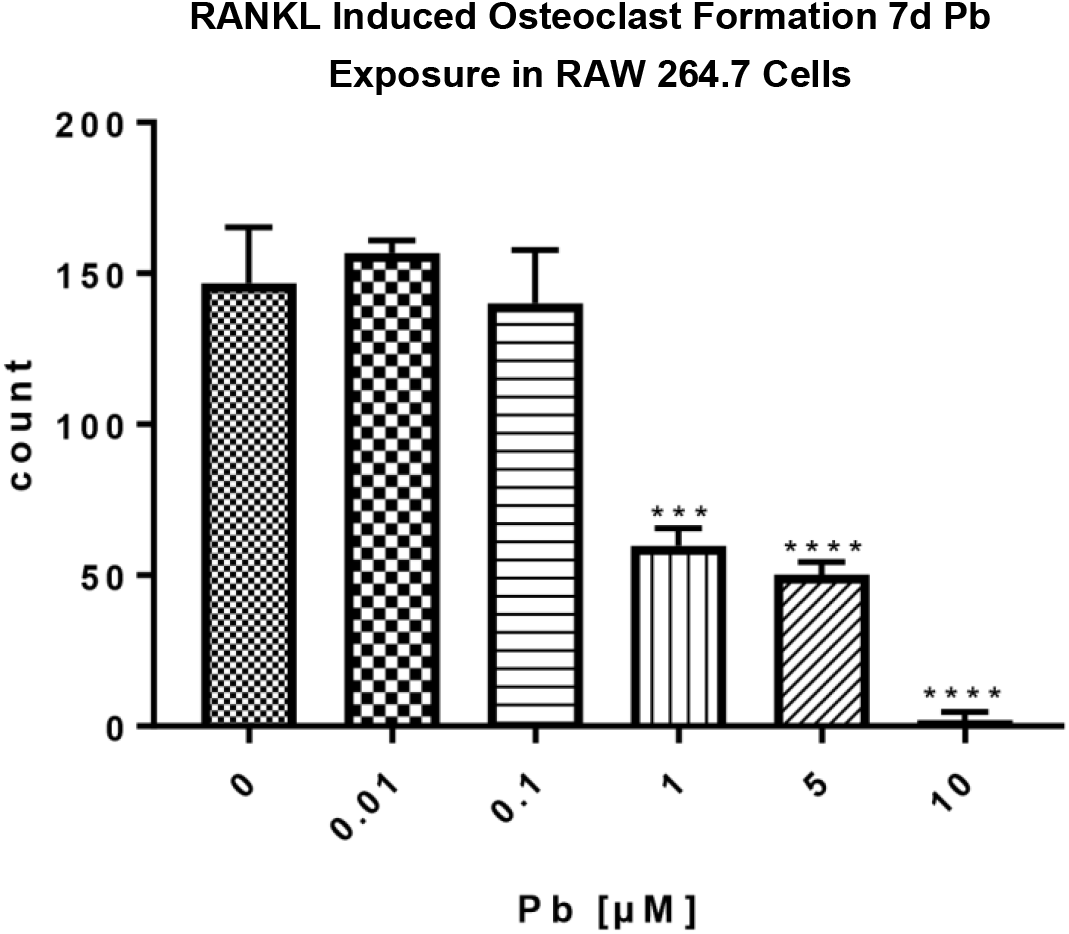
Pb induced inhibition of osteoclast formation scored by counting under phase contrast light microscopy. Unpaired *t* test was performed to determine statistical significance (n = 3) ***p ≤ 0.001. ****p ≤ 0.0001.

### GM-CSF Production

To clarify the identity of the newly formed cells in RAW 264.7 cells induced by continuous exposure to 5.0 and 10 µM Pb, GM-CSF production was analyzed. GM-CSF regulates the adaptive immune system through regulation of T-cell differentiation by the action of DC’s (Bhattacharya P. et al 2015). It also inhibits osteoclast formation and stimulates DC formation (33). Long-term exposure to Pb only, did not alter the production of GM-CSF in RAW 264.7 cells. However, stimulation by RANKL revealed a trend in which Pb dose-dependently increased GM-CSF production (Figure 5). This trend corresponded with the presence of non-adherent dendritic-clusters in culture. It is not clear whether the production of GM-CSF here precedes DC differentiation or is a consequence of DC formation.

**Figure 5.**
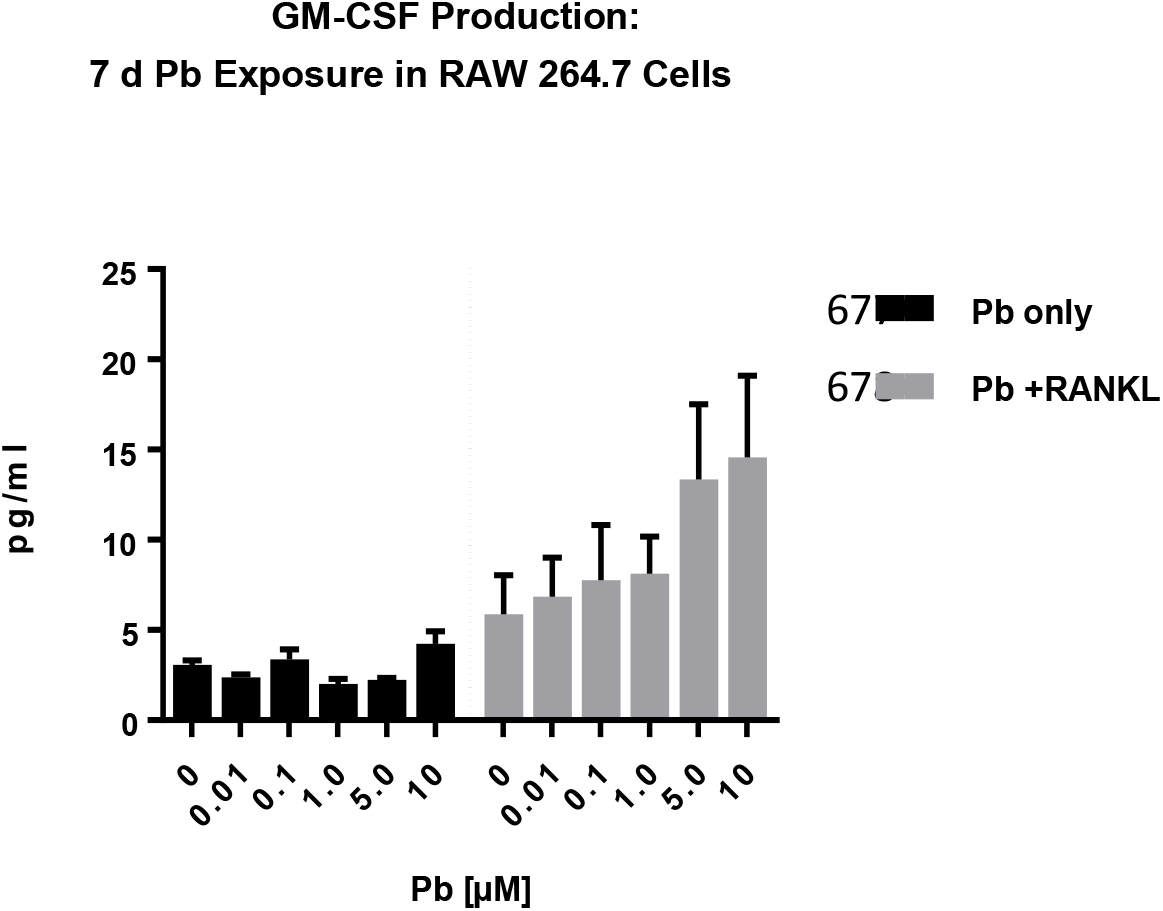
A trend of Pb induced dose-dependent increase in GM-CSF production in RAW 264. 7 cells exposed to Pb and stimulated with RANKL for 7 d was present (n = 3). Pb alone did not increase GM-CSF production above base-line (n = 3). Two-way ANOVA and Unpaired *t* test were performed to determine statistical significance. Experiments were replicated twice.

### NFκB Activity

NFκB activity is essential to osteoclastogenesis (34). To validate inhibition of osteoclastogenesis and to elucidate a possible mechanism of action, NFκB activity was examined in RAW Blue cells. A dose-dependent inhibition of NFκB activity by Pb was seen with 5 d exposures.

Significant inhibition of NFκB occurred between 1.0 and 10 µM Pb, and as low as 0.1 µM Pb with co-exposure to RANKL (Figure 6).

**Figure 6.**
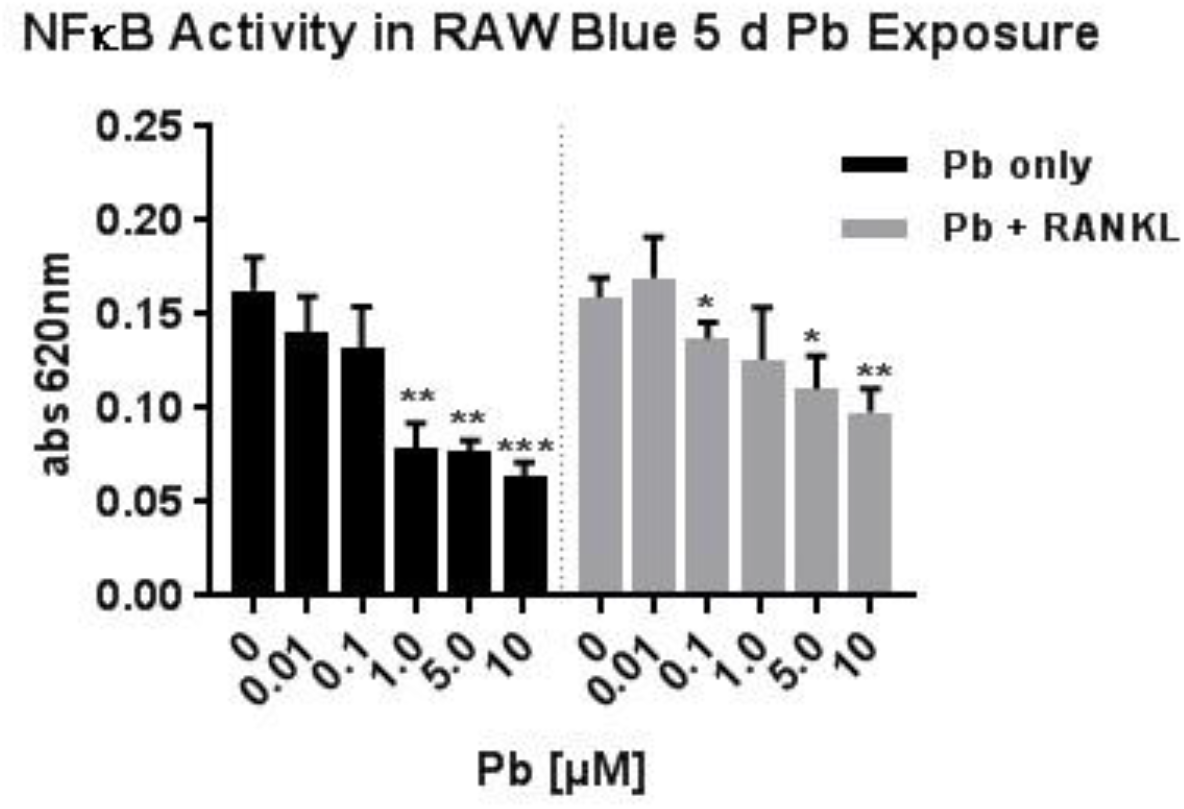
Pb inhibited NFkB activity in RAW Blue cells with 5 d Pb exposure (n = 3) with and without RANKL stimulation. Activity decreased in a dose-dependent manner. Unpaired *t* test was performed to determine statistical significance. *p <0.05, **p ≤ 0.01, ***p≤ 0.001.

### IL-1α production

IL-1α is also involved in differentiation and multinucleation of monocyte/macrophage cells. It was reported to have the potential to induce osteoclastogenesis via a RANK/RANKL independent mechanism, though it does co-operate with RANKL induced osteoclast formation (35). To further validate that long-term exposure to Pb inhibits the differentiation of osteoclasts, IL-1α was examined. Exposure to Pb alone or in conjunction with RANKL for 7 d revealed significant decrease in the production of IL-1α by RAW 264.7 cells at all concentrations, compared to control (Figure 7).

**Figure 7.**
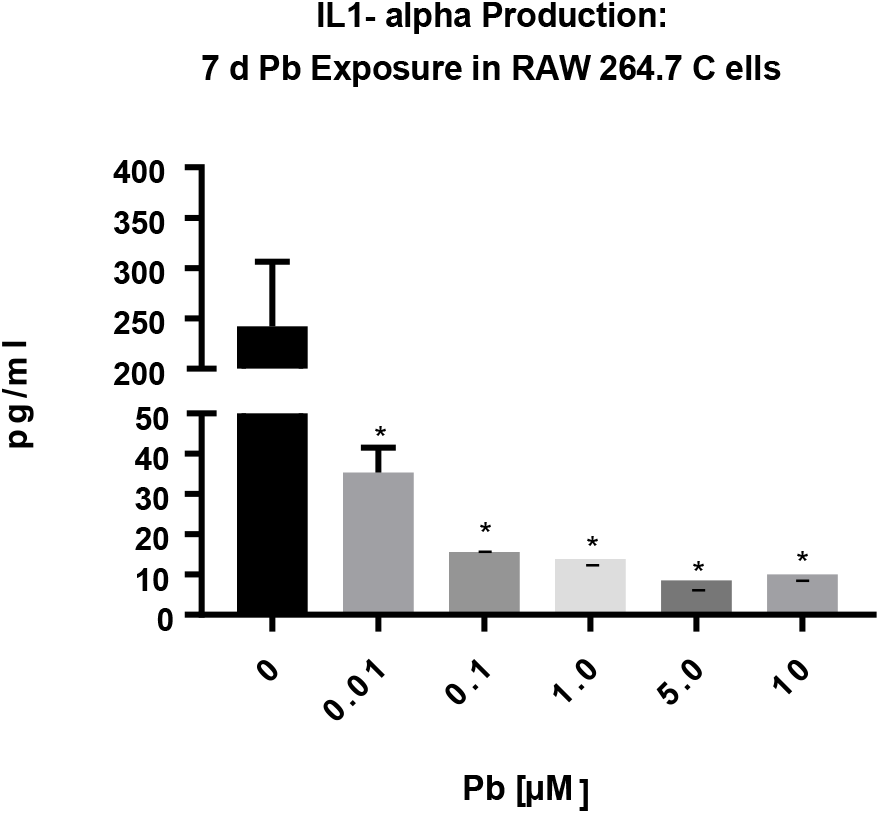
Long-term exposure to Pb inhibited the production of IL-1α, a potent inflammatory cytokine involved with osteoclast function and formation. Unpaired *t* test was performed to determine statistical (n = 3). Error bars represent standard error of the mean, * p < 0.05.

### p38/MAPK Activity

p38/MAPK plays an important role in cell survival, proliferation, inflammatory response, and differentiation (36). It functions downstream of RANK/RANKL and TNF-α/TNFR stimulation. Inhibition of p38 blocks RANKL and TNF-α induced osteoclastogenesis (35). Changes to p38/MAPK activity occur early, and therefore the effect of Pb on its activity was examined at 24 h. Pb inhibited p38/MAPK activity in a dose-dependent manner in RAW 264.7 cells. Pb alone and Co-exposure of Pb with RANKL resulted in prominent decrease in p38/MAPK activity. Significant inhibition occurred at 0.1, 5.0, and 10 µM Pb (figure 8A) and between 1.0 and 10 µM Pb + RANKL (figure 8B), respectively.

**Figure 8.**
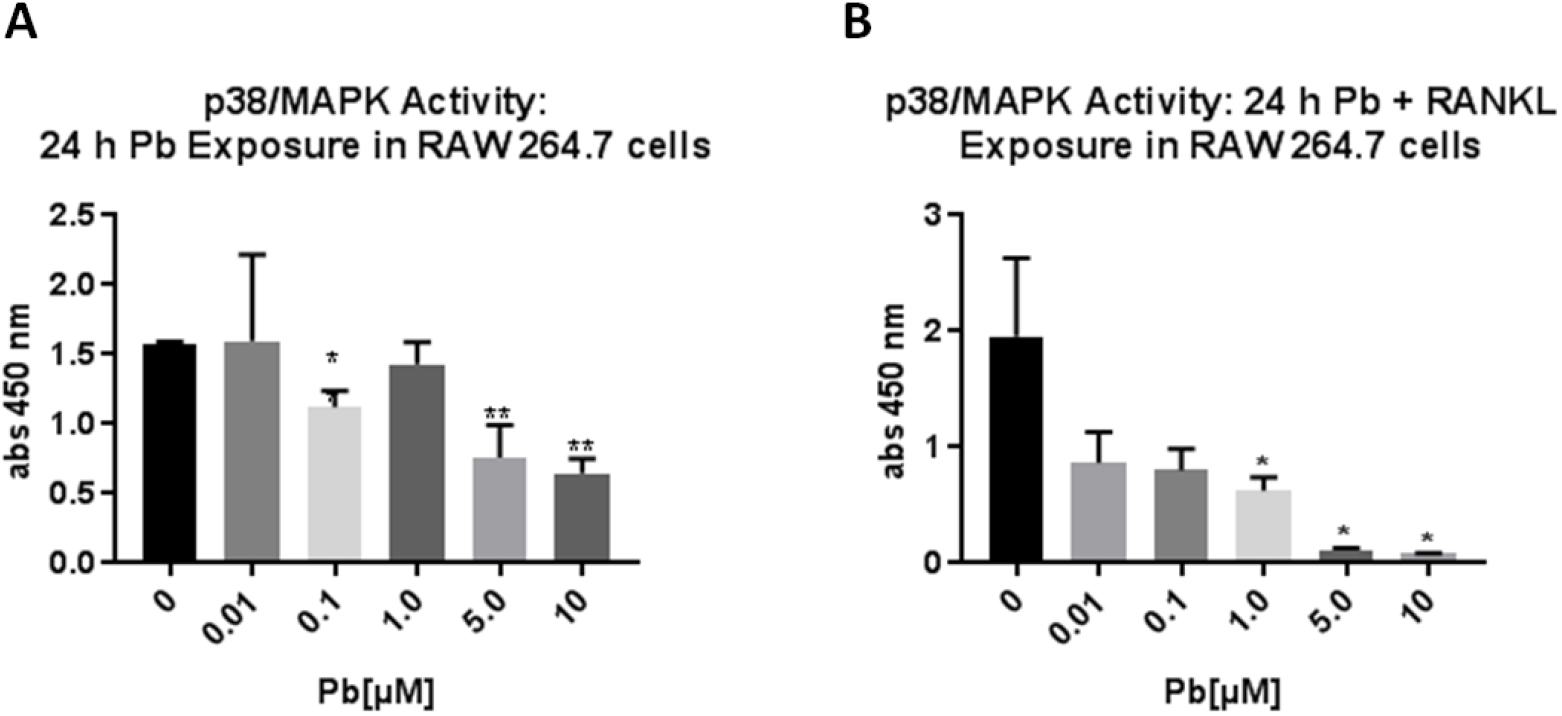
Effect of Pb on p38/MAPK activity in RAW 264.7 cells. Pb inhibited p38MAPK activity in a dose-dependent manner with (A) and without RANKL (B). Unpaired *t* test and one-way ANOVA were performed to determine statistical significance (n = 3). Experiments were replicated twice * p <0.05, **p ≤ 0.01.

### Immunofluorescent staining

#### CD11c vs CD11b and DEC205 in RAW 264.7 cells

Indeed, when RAW 264.7 cells were fluorescently stained with antibody markers of DC, CD11c and DEC205, a dose-dependent population shift towards dendritic phenotype was observed with long-term exposure to Pb. RAW 2647 cells exposed to Pb only or Pb and RANKL developed a population of cells of various sizes and high expression of cell surface markers CD11c vs.

CD11b (Figure 9) and DEC 205 (Figure 10). This occurred in a dose-dependent manner, with greatest expression in cells exposed to concentrations of Pb ranging from 1 to 10 µM. CD11c^+hi^/CD11b^+hi^ cells are reported to be immature DC while CD11c^+hi^/CD11b^+low^ are indicative of mature DC. Additionally, DEC205 is involved with antigen presentation and increased expression is associated with mature DC (37). These results validate that non-adherent clustered cells formed by long term (5 to 7 d) exposure to high concentration (5.0 and 10 µM) of Pb are DC’s.

**Figure 9.**
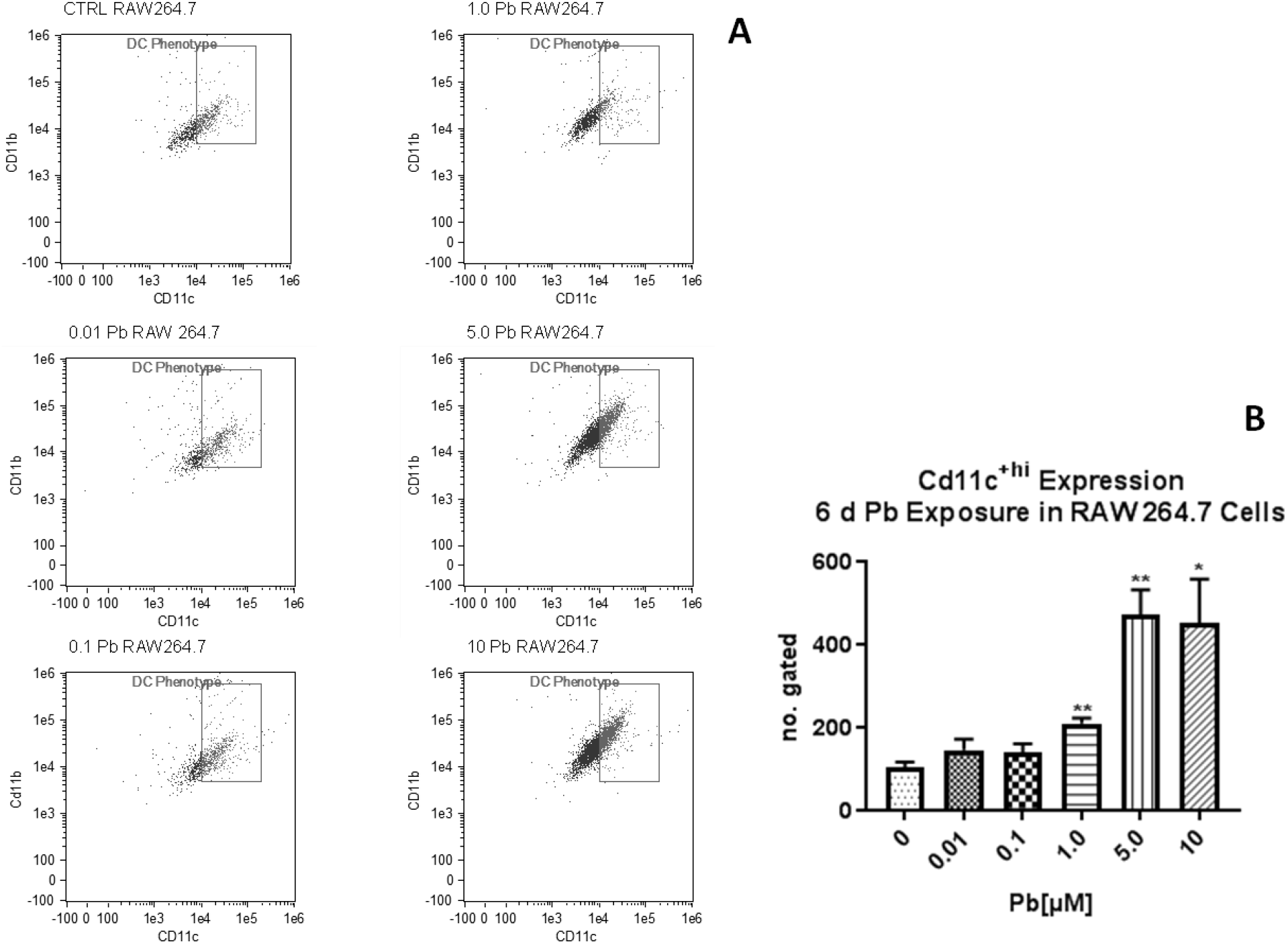
Pb induced dendritic phenotype in RAW 264.7 cells. Scatter plots are representative and reflect CD11b vs CD11c expression as determined by intensity (10,000 objects total), in RAW 264.7 cells exposed to Pb for 6 d; images are representative (A). CD11c+/CD11b+ cells are reported to be immature DC., Pb induced a dose-dependent increase in cells reflective immature DC, 3000 objects were counted and gated for high intensity of CD11c vs CD11b, mean value of gated area is represented in bargraph (B). Unpaired *t* test was performed to determine statistical significance, and error bars reflect standard error of the mean (n = 3), * p <0.05, **p ≤ 0.01.

**Figure 10.**
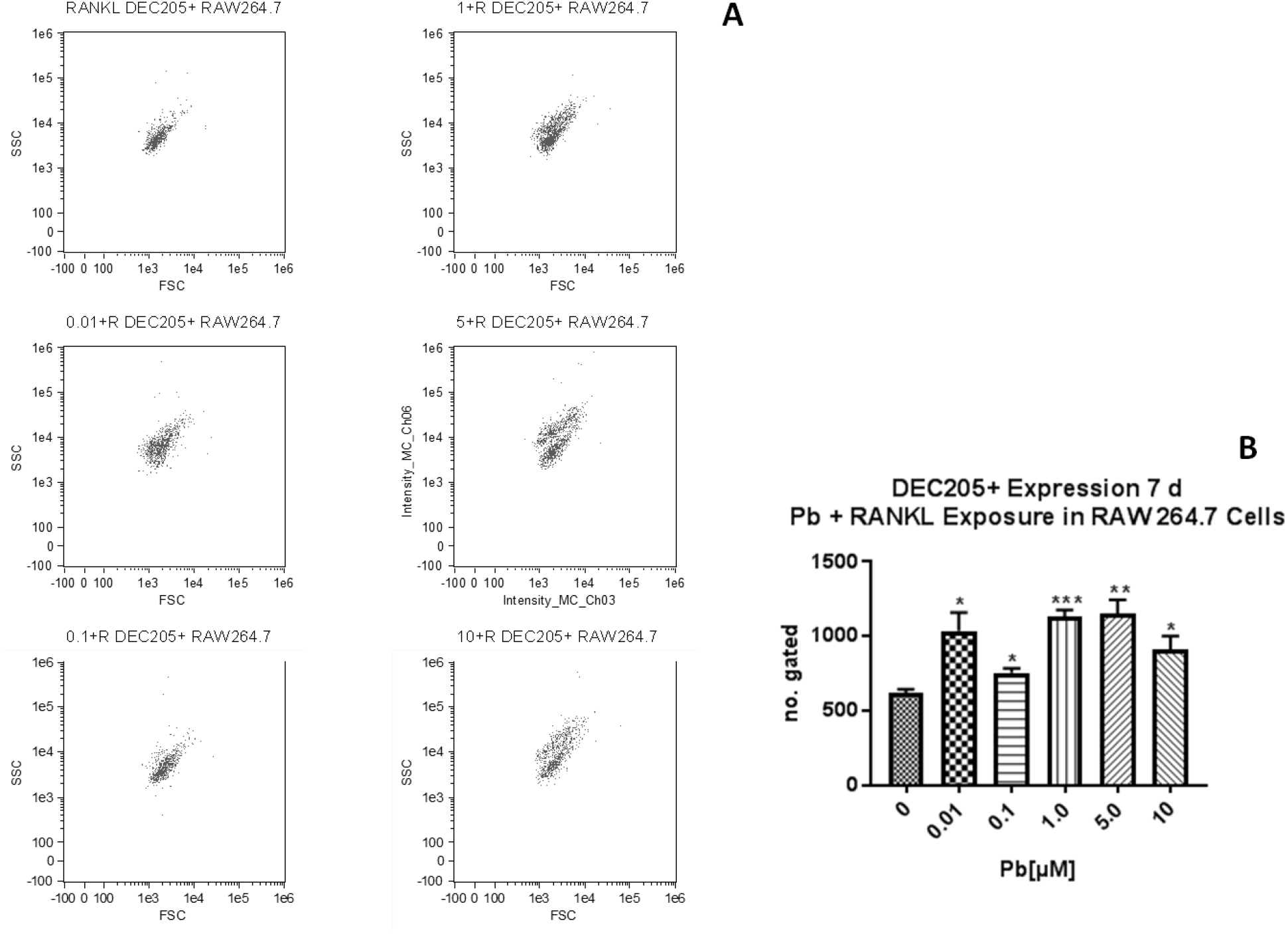
Pb and RANKL co-exposure (7 d) induced the differentiation of mature DC in monocyte/macrophage RAW 264.7 cells as evidenced by DEC205^+^ cell population shift. 3000 objects were analyzed and single cells with high intensity of DEC205 were gated, and representative Images are displayed (A). Gated population was counted and mean values expressed by bar graph. Unpaired *t* test was performed to determine statistical significance (n =3). Error bars reflect standard error of the mean. * p <0.05, **p ≤ 0.01, p≤0.001.

#### Phagocytic Population

Monocyte/macrophage cells exist in a heterogeneous population (37, 38). As such, long-term exposure to Pb in RAW 264.7 cells did not produce differentiation of all cells, but resulted in a mixed population of cells consisting of monocyte/macrophage, osteoclast, nonadherent activated monocytes, and DC. However, Pb dose-dependently shifted the predominant population of cells present in culture. This was evidenced by phagocytic capacity. Exposure to 1 µM Pb for 7d resulted in the appearance of non-adherent monocyte/macrophage cells that were highly phagocytic compared to control (Figure 11). Pb not only induced differentiation to DC, but stimulated a highly phagocytic monocyte/macrophage population *in vitro*.

**Figure 11.**
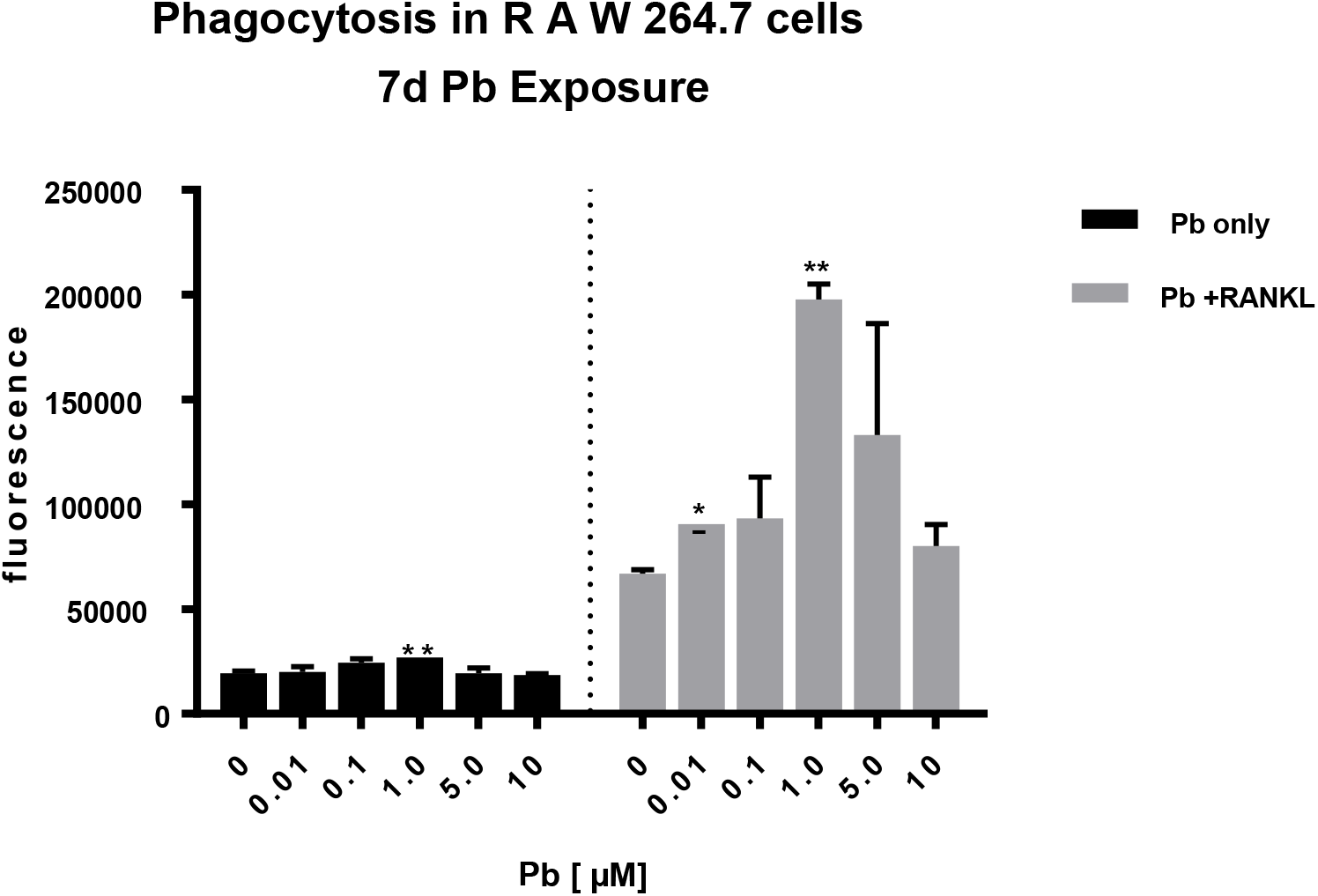
Long-term exposure to Pb in RAW 264.7 cells induced a highly phagocytic cell population that was dramatically enhanced by RANKL stimulation. Unpaired *t* test was performed to determine statistical significance, and error bars reflect standard error of the mean (n = 3). * p <0.05, **p ≤ 0.01.

#### Differentiation in BMMØ

To determine whether the effect of Pb on differentiation in RAW264.7 cells was relevant to monocyte/macrophage cells in general and not a result of cell-line characteristics, Pb was tested in rat bone-marrow monocyte/macrophage (BMMØ) -- BMMØ were derived from rat bone-marrow cells treated with M-CSF. Osteoclastogenesis was induced by culturing BMMØ on tissue-culture treated plates (without requirement of RANKL). Long-term Pb exposure inhibited osteoclast formation in rat BMMØ. Osteoclasts formed only in control, 0.01 µM, and 0.1 µM exposure groups; with osteoclast formation declining from 0 to 0.1 µM Pb exposure, consecutively (figure 12). Furthermore, osteoclasts did not form in cells exposed to Pb ranging in concentration from 1. 0 to 10 µM, instead cells became amoeboid and did not attach to the plate surface (images not shown). Interestingly, as mentioned above, attachment to tissue-culture treated surface triggered osteoclast formation in BMMØ. Furthermore, this phenomenon of bifurcation in characteristic of monocyte/macrophage cells was consistent with the effects seen in RAW 264.7 cells with long-term Pb exposure at the corresponding concentrations (1.0 to 10 µM Pb). In which, non-adhesive amoeboid cells gave rise to phagocytic monocyte/macrophage and dendritic-like cells. Therefore, cells were probed for CD11c expression; a cell-surface marker for dendritic cells.

**Figure 12.**
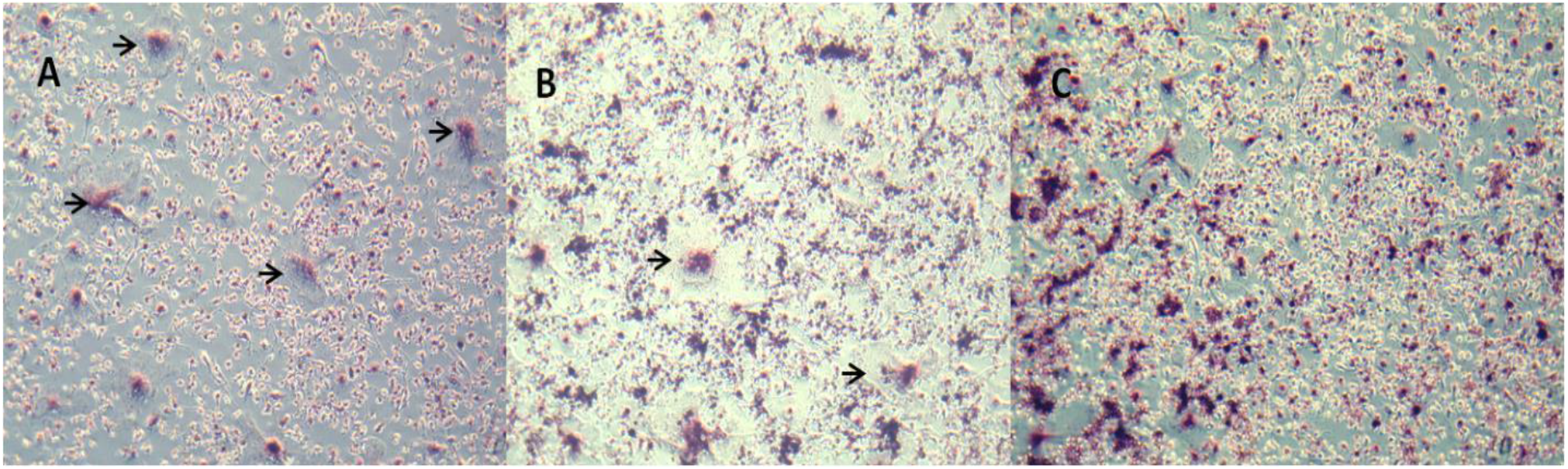
Osteoclastogensis in BMMØ exposed to Pb for 7 d in vitro. Long-term Pb exposure in BMMØ resulted in a dose-dependent inhibition of osteoclast formation. Osteoclasts formed in control (A) 0.01 µM Pb (B) and 0.1 µM Pb (C) only. Osteoclasts did not form in BMMØ exposed to concentrations of Pb ranging from 1.0 to 10 µM Pb. Images are representative of fixed TRAP-stained cells and were taken via phase contrast light microscopy (n = 3). *p <0.05,

Immunofluorescent antibody staining for CD11c revealed Pb significantly induced the formation of a CD11c^+ hi^ population in rat BMMØ exposed to 0.1 to 10 µM concentrations for 9 d (Figure 13). In response to 9 d Pb exposure, a trend of increasing GM-CSF production was observed, with significant increase in BMMØ exposed to 0.1 and 5 µM Pb (figure 14). However, GM-CSF production declined again at 10 µM Pb. The results of long-term exposure to Pb in BMMØ derived from rat bone-marrow cells were consistent with changes in osteoclastogenesis and the induction of DC formation seen in RAW 264.7 cells.

**Figure 13.**
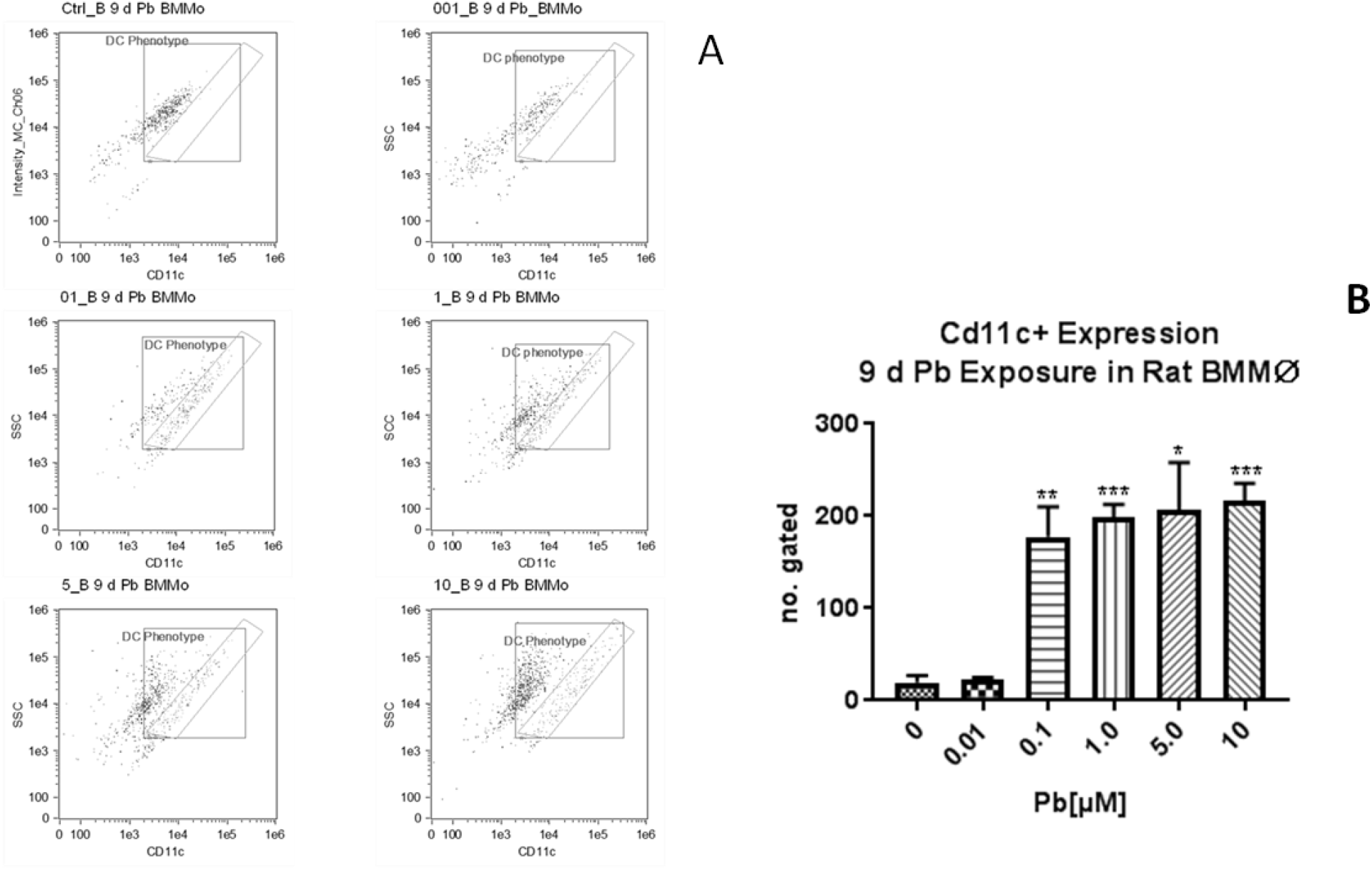
9 d Pb exposure significantly induced a dendritic population in BMMØ cells at concentrations ranging from 0.1 to 10 µM. Flow cytometry images are representative of CD11c^+^ dose-dependent induction (dendritic phenotype) in BMMØ by long-term Pb exposure (A). 3000 objects were analyzed and population gating of single cells with CD11c^+hi^ expression was performed; mean values are represented by bar graph (B). Statistical analysis was determined using unpaired t test (n =3). *p <0.05, **p ≤ 0.01, ***p≤ 0.001.

**Figure 14.**
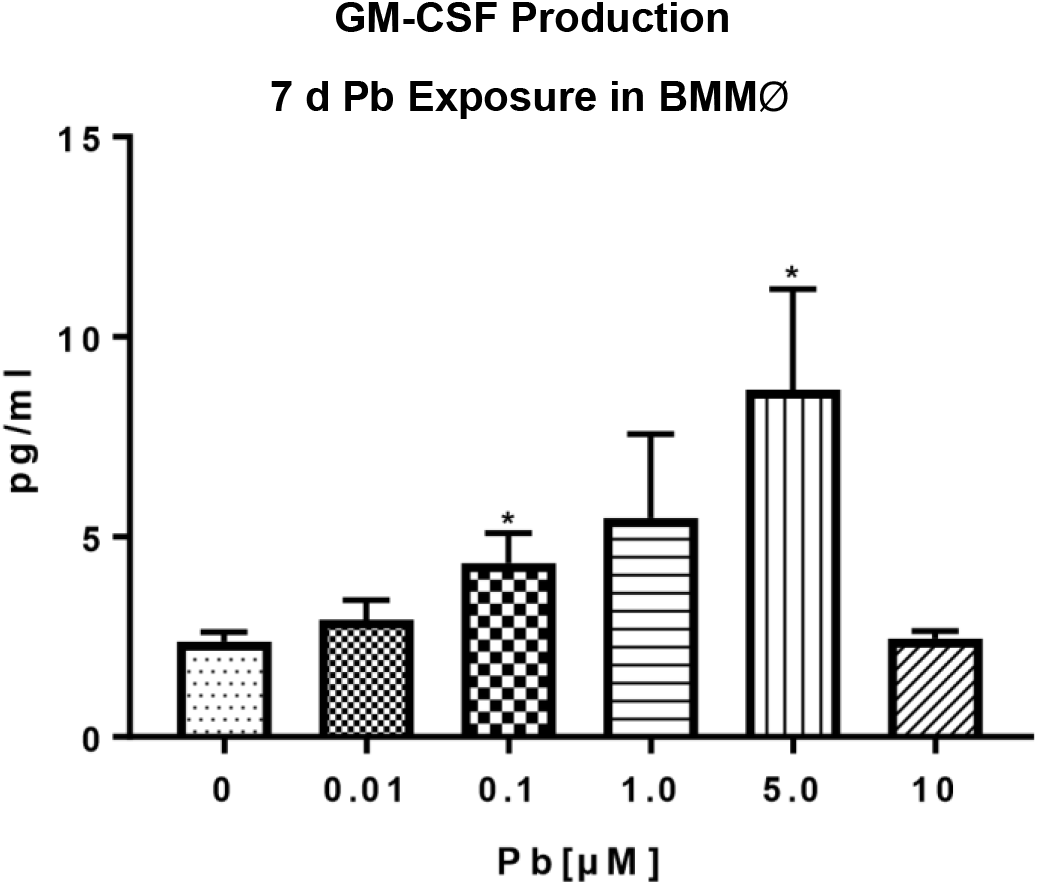
In response to long-term Pb exposure, a trend of increasing GM-CSF production was observed in rat BMMØ as assayed by ELISA. Production was significantly increased at 0.1 and 5.0 µM Pb. Though increased production was not detected with exposure to 10 µM Pb, these results correspond with the effect of pro-longed Pb exposure on GM-CSF production seen in RAW 264.7 cells (figure 5) stimulated with RANKL. Statistical analysis was determined using ANOVA and unpaired t test (n = 6). *p 0.05.

## Discussion

### Cell Survival and Differentiation

Long-term exposure of monocyte/macrophage cells to Pb was not associated with cell death or inflammation, but differentiation. Pb induced the differentiation of DC cells in murine RAW 264.7 monocyte/macrophage cell-line and rat BMMØ cells through a process involving p38/MAPK inhibition and GM-CSF production. Inhibition of p38/MAPK is an early event and likely responsible for the reduction in TNF-α production and subsequently proliferation (39), as its activation is involved with TNF-α biosynthesis (40).

Further examination of Pb’s ability to affect differentiation revealed a bifurcation in osteoclast and DC formation from monocyte/macrophage cells. When examined in an osteoclast model of differentiation, Pb produced a dose-dependent alteration in osteoclastogenesis; inhibiting osteoclast formation with increased concentrations. Osteoclast inhibition coincided with DC formation, and a third population of highly phagocytic activated monocyte/macrophage was also produced. Osteoclastogenesis takes place in three stages: activation, fusion, and polarization – without fusion osteoclasts do not form. The nonadherent phagocytic population formed at the mid-range concentration of Pb, was likely a consequence of failure in the fusion stage of osteoclastogenesis.

Additionally, though we had observed dendritic-clusters and large dendritic-like cells with Pb only exposure, we did not detect an appreciable presence of GM-CSF, the cytokine necessary for DC production. However, in the presence of RANKL, we did detect rising amounts of GM-CSF, in-line with increases in Pb concentrations. It is possible that RANKL results in the maturation of the dendritic-clusters/Dendritic-like cells produced by Pb only exposures. Pb only cells may be immature or precursor DC’s. Detectable levels of GM-CSF present with co-exposure of Pb + RANKL may be a by-product of GM-CSF receptor shedding from mature DCs or DC activation state.

### Pb Inhibition of Osteoclastogenesis

While Pb alone could induce DC formation, RANKL was required for the differentiation of osteoclasts from monocyte/macrophage cells. RANKL activates several transcription factors involved with cell differentiation and osteoclastogenesis: NFκB, p38/MAPK, c-Fos, NFATC1, and PU.1 (41), several of which were found to be altered by long-term exposure to Pb in a dose-dependent manner. NFκB activation is essential for osteoclast formation, and TNF-α co-operates with M-SCF and RANKL to respectively induce and enhance osteoclastogenesis. Pb’s inhibition of osteoclastogenesis in RAW 264.7 cells were associated with a dose-dependent inhibition of NFκB and p38/MAPK, and TNF-α production. Additionally, Pb similarly inhibited IL-1 production. IL-1 is reported to work synergistically with RANKL, but can induce osteoclastogenesis through a RANK/RANKL independent mechanism involving the activation of NFκB and p38 (35). Therefore, it is concluded that Pb inhibits osteoclastogenesis by inactivation of the p38/MAPK pathway, and through a mechanism involving the inactivation of NFκB and inhibition of IL-1α production.

Pb’s inhibition of key components to osteoclastogenesis: NFκB and p38/MAPK, and of TNF-α and IL-1α coincided with induction of DC differentiation. The activation state of p38/MAPK has been identified as playing a pivotal role in the formation of both osteoclasts and DC, with activation necessary for osteoclastogenesis, and inhibition favoring DC formation (42).

Activated MAPKs regulate the expression of genes involved in differentiation and osteoclastogenesis through phosphorylation of downstream transcription factors (Yokoyama et al, 2013), such as c-Fos. Pb may alter the expression of c-Fos, which is a transcription factor involved in the regulation of NFATc, and responsible for cell-cell fusion in osteoclatogenesis. p38/MAPK controls the gene expression of transcription factors through phosphorylation and cross-talk with other signaling pathways (43). c-Fos gradually accumulates during osetoclastogenesis, and induction of its expression subsequently leads to the activation of NFATc1, which is involved with cell-cell fusion (44)

The expression of c-Fos is regulated by MAPK and PKC pathways (45). It is important to note that Pb did not significantly alter PKC activity in RAW 264.7 cells (data not shown), which can activate p38/MAPK. Intact PKC activity may then be responsible for continued cell survival, while it is through the action on p38/MAPK activity that Pb alters differentiation in monocyte/macrophage-lineage cells, with p38/MAPK activation or inactivate functioning as a switch between monocyte/macrophage differentiation to osteoclasts or dendritic cells (44).

### Pb Induced DC Formation

Additionally, Pb induction of DC formation was associated with GM-CSF production. Dysregulation of GM-CSF was reported to play a possible role in the infiltration of the CNS by phagocytic monocytes leading to tissue injury and neurological deficits (46). In the presence of RANKL, long-term Pb exposure induced the production of GM-CSF in monocyte/macrophage cells, in vitro, as Pb concentrations increased. GM-CSF is a growth factor commonly used in the generation of DC *in vitro*, and depending on concentration can induce both immature and mature DC in BMC without additional stimuli (47). Long-term Pb in monocyte/macrophage cells resulted in the spontaneous formation of both immature and mature DC, and this may be a function of increased GM-CSF production though it is not clear whether GM-CSF present in culture was a prerequisite or consequence of DC formation. Detected GM-CSF in the supernatant of monocyte/macrophage cells cultured long-term in the presence of Pb may have resulted from the upregulation and subsequent shedding of the GM-CSF receptor from cell surfaces. RAW 264.7 cells constitutively shed their M-CSF receptors which then function as the soluble cytokine (27), and the same may be true for GM-CSF. M-CSF and GM-CSF are reported to have opposed functions in the differentiation of monocyte/macrophage cells (48). Furthermore, Pb may be capable of altering the expression of genes, such as PU.1, which are involved in regulating the expression of monocyte/macrophage cell-surface receptors M-CSF and GMCSF (49, 50).

PU.1 is a transcription factor involved with the proliferation and differentiation of monocyte precursor cells to granulocytes and macrophage, it regulates gene transcription of *c-fos* which determines the responsiveness of myeloid progenitors to M-CSF or GM-CSF (22, 49). Prolonged high expression of PU.1 sensitizes monocyte/macrophage lineage cells to GM-CSF stimulation and promotes granulocyte and DC formation (51). Therefore, Pb may increase GM-CSF production and induced DC formation through upregulation of transcription factor PU.1. Further investigation is being conducted by our lab to determine the involvement of c-Fos and PU.1 with the phenomenon described.

## Conclusion

Pb induces differentiation of DC and inhibition of osteoclastogenesis in monocyte/macrophage cells by directly inhibiting the p38/MAPK pathway. Pb’s ability to alter the population of immune cells of monocyte/macrophage lineage by the dose-dependent inhibition and induction of osteoclasts, and activated macrophage/monocytes and dendritic cells respectively, indicates that Pb can affect both the innate and adaptive immune system. Furthermore, in the absence of LPS and ATP, Pb inhibited inflammation: Pb inhibited the production of inflammatory mediator IL-1α, which is also involved in cell death (necrosis, apoptosis, and pyroptosis) (52), and TNF-α and NFκB, which play integral roles in the generation of inflammation and monocyte maturation and differentiation. Interestingly, it has been reported that PU.1 deficient mice showed attenuation of TNF-α and NFκB secretion from mature macrophage when challenged with LPS (22). This highlights a possible an involvement of PU.1 gene expression in the mechanism of action of Pb on monocyte/macrophage differentiation (investigation by our lab is being conducted). Though Pb stimulated increased production of GM-CSF, GM-CSF can exacerbate or ameliorate numerous inflammatory related conditions (47, 53). Moreover, Pb induction of GM-CSF may contribute to the formation of both immature and mature DC, which have differential effects on the induction of T-cells (54, 55).

Therefore, it is concluded that Pb functions as an immune-modulator; by inducing differentiation of monocyte/macrophage cells to DC in lieu of proliferation and multinucleation of macrophage, and attenuating the production of inflammatory mediators. These findings have important implications for the role of Pb in the genesis and progression of a multitude of immune-mediated diseases, such as bone disorders (56), multiple sclerosis (57), rheumatoid arthritis (58), and human immunodeficiency virus infection (59). Further studies conducted in vivo are required.

## Acknowledgements

The department of Pharmaceutical Science, in the College of Pharmacy and Health Sciences at St. John’s University for funding this research.

